# Differential assembly of RNP granules via activation of distinct dsRNA sensors by adenovirus mutants

**DOI:** 10.64898/2026.02.17.706363

**Authors:** Robert T. Steinbock, Orlando B. Scudero, Joseph M. Dybas, Amber R. N. Abbott, Katarzyna Kulej, Richard Lauman, Holly Chan, Eva L. Agostino, Namrata Kumar, Skyler Briggs, Nicholas A. Parenti, Yize Li, James M. Burke, Susan R. Weiss, Alexander M. Price, Matthew D. Weitzman

## Abstract

Recognition of dsRNA triggers antiviral defense mediated by PKR and OAS3/RNase L pathways through translational arrest and RNA decay. This is accompanied by assembly of distinct cytoplasmic ribonucleoprotein (RNP) condensates termed stress granules (SGs) and RNase L-dependent bodies (RLBs). Here we show that adenovirus infection differentially modulates dsRNA sensors and RNP granule assembly. Infection with splicing-defective ΔE4 mutant leads to dsRNA accumulation and activation of both PKR and OAS3/RNase L, promoting formation of RLB-like granules. In contrast, mutants lacking virus-associated (VA) RNAs trigger PKR activation and assembly of SGs despite absence of detectable dsRNA. Proteomic analysis revealed distinct protein compositions of canonical SGs and RLBs, which were reflected in virus-induced granules. While ΔVA-induced granules were PKR-dependent, ΔE4 mutants induced RLB-like granules independently of PKR and RNase L. In these cells, granule assembly coincided with translational arrest independent of eIF2α phosphorylation, indicating additional pathways linking nuclear dsRNA sensing to translational control and RNP granule assembly during viral infection. These findings provide novel insights into how distinct dsRNA sensors modulate translation and RNP condensates in response to stress.

## INTRODUCTION

Among the numerous strategies used by cells to counter viral infections, recognition of double stranded RNA (dsRNA) is an essential host defense mechanism (Hur, 2019; Chen & Hur, 2022; Cottrell *et al*, 2024). In this context, three major signaling pathways are regulated by cellular dsRNA sensors. These include RIG-I-like receptors (RLRs), Protein Kinase RNA-activated (PKR), and oligoadenylate synthetase 1-3/RNase L (OAS1-3/RL) (Chan & Gack, 2016; Schlee & Hartmann, 2016). Activation of RLRs (e.g. MDA5 and RIG-I) triggers multimerization of the mitochondrial outer membrane protein MAVS, which in turn recruits and activates various signaling molecules (Yoneyama *et al*, 2004; Kato *et al*, 2006; Gitlin *et al*, 2006; Kato *et al*, 2011). This leads to phosphorylation of the transcription factors IRF3 and IRF7, with subsequent induction of interferons (IFNs) and interferon-stimulated genes (ISGs) (Fitzgerald *et al*, 2003; Rehwinkel & Gack, 2020; Onomoto *et al*, 2021). PKR is activated upon binding to dsRNA, which drives its sequential dimerization and autophosphorylation at residues T446 and T451 (Dey *et al*, 2005; Dauber & Wolff, 2009; Zhang *et al*, 2001). Activated PKR then phosphorylates eIF2α at S51, blocking translation initiation of bulk cellular mRNAs (McCormick & Khaperskyy, 2017; Pakos-Zebrucka *et al*, 2016). Upon dsRNA binding, OAS1-3 enzymes synthesize the secondary metabolite 2’-5’ oligoadenylate (2’-5’A), which activates latent RNase L (Hovanessian *et al*, 1979; Silverman *et al*, 1983; Silverman, 2007). Activated RNase L then promiscuously degrades both viral and host RNAs, promoting widespread cytoplasmic mRNA turnover and translational arrest (Donovan *et al*, 2017; Burke *et al*, 2019). Activation of these pathways illustrates how cells employ different mechanisms in response to dsRNA, which leads to distinct downstream effects.

An additional outcome of activation of PKR and RNase L pathways is the assembly of distinct cytoplasmic ribonucleoprotein (RNP) condensates, namely stress granules (SGs) and RNase L-dependent bodies (RLBs) (McCormick & Khaperskyy, 2017; Burke *et al*, 2020; Watkins & Burke, 2024a; Pakos-Zebrucka *et al*, 2016). Both granules co-stain for G3BP1 and PABPC1, however, RLBs are distinguished by their smaller size, more spherical morphology, and the bulk relocalization of PABPC1 to the nucleus due to widespread RNase L-dependent degradation of cytosolic mRNA (Burke *et al*, 2020, 2019). Among their functions, SGs play a more general role in blocking translation and sequestering RNA and proteins to mitigate cellular stress (Eiermann *et al*, 2020; Protter & Parker, 2016). SGs are also thought to concentrate viral and host RNA sensors, thereby enhancing innate antiviral signaling pathways (Reineke *et al*, 2015; Onomoto *et al*, 2014; Paget *et al*, 2023), although recent studies suggest this model may not apply across all contexts (Burke *et al*, 2024; Watkins & Burke, 2024a; Corbet *et al*, 2021, 2022; Langereis *et al*, 2013). Diverse viruses have been demonstrated to facilitate infection by either inhibiting SG assembly or interfering with their functions by co-opting host proteins (Zhang *et al*, 2019; Brownsword & Locker, 2023; Eiermann *et al*, 2020). In contrast, RLBs are thought to accumulate RNAs cleaved by RNase L, assisting the cellular RNA decay machinery (Watkins & Burke, 2024b; Burke *et al*, 2020). This effect has been reported to inhibit infection by several flaviviruses, suggesting that RLBs contribute to host antiviral defenses (Watkins & Burke, 2024a, 2024b; Burke, 2023).

Adenovirus (AdV) is a well-studied, linear double-stranded DNA virus which has been involved in seminal discoveries in RNA biology and virus-host interactions. AdV possesses a compact genome (∼36 kbp) organized into several transcriptional units on both the top and bottom strands (Berk AJ *et al*, 2013). Transcription from these units is regulated by alternative splicing, generating a wide array of viral transcripts throughout infection (Price *et al*, 2022b; Donovan-Banfield *et al*, 2020; Alemán *et al*, 2022). This bidirectional architecture has been speculated to produce intermolecular viral dsRNAs formed by mirrored exons and introns from opposing strands, with potential for being detected by antiviral dsRNA sensors (Jacobs & Langland, 1996). Efficient processing of viral pre-mRNAs is regulated by the viral E1B-55K/E4orf6 complex which harnesses a cellular E3-ubiquitin ligase (Herrmann *et al*, 2020), and we recently discovered that splicing-defective mutants lacking these proteins accumulate abundant nuclear dsRNA which activates PKR (Spurgeon & Ornelles, 2009; Price *et al*, 2022a). All human adenoviruses also encode one or two copies of highly structured non-coding virus-associated RNAs (VA RNA I and II) that counter PKR activation (Ma & Mathews, 1996; Punga *et al*, 2020; Vachon & Conn, 2016). Pioneering studies have shown that infection with viral mutants lacking VA RNAs induces PKR phosphorylation, leading to translation shutoff and impaired late gene expression (Bhat & Thimmappaya, 1984; Schneider *et al*, 1985; Kitajewski *et al*, 1986). These findings contributed to the identification of VA RNA I as a PKR antagonist, leading to the presumption that dsRNA is formed during adenovirus infection (Mellits *et al*, 1990; Wahid *et al*, 2009; Wilson *et al*, 2014). However, subsequent studies have failed to detect dsRNA in cells infected by wild-type or VA-deficient viruses, suggesting PKR is activated by another distinct mechanism during infection with ΔVA mutants (Price *et al*, 2022a; Maran & Mathews, 1988).

Here we examined how infection with AdV serotype 5 (Ad5) variants can modulate distinct dsRNA sensors and impacts assembly of RNP granules. Comparing mutants that produce dsRNA (ΔE4, *dl1004*) (Bridge & Ketner, 1989) with mutants that do not (ΔVA, *dl-sub720*) (Bhat & Thimmappaya, 1984) revealed that viral dsRNA formed during ΔE4 infection activates both PKR and OAS3/RNase L pathways, leading to formation of granules similar to RLBs. Conversely, infection with the ΔVA mutant does not yield detectable dsRNA but still activates PKR, resulting in PKR-dependent assembly of stress granules. Using APEX-proximity labeling during treatment with chemical stressors (Markmiller *et al*, 2018), we identified proteins differentially enriched in RNP granules induced by sodium arsenite (SA) or during transfection with the dsRNA analog polyinosinic:polycytidylic acid, poly(I:C). Characterization of granules formed in infected cells using immunofluorescence demonstrated that virus-induced granules are morphologically and compositionally analogous to their chemically-induced counterparts. Infection of cells lacking dsRNA sensors showed that infection with the ΔE4 mutant promotes translational arrest and assembly of cytoplasmic granules with properties similar to canonical RLBs, independent of both PKR and RNase L. These results suggest that alternative or redundant pathways may regulate translation and RNP granule induction in response to viral infection or accumulation of nuclear dsRNA. Our study provides novel insights into the formation and regulation of RNP complexes during virus infection, expanding our understanding of dynamic host-virus interactions and opening new avenues for investigating cytoplasmic RNP condensates.

## RESULTS

### Distinct dsRNA sensors are activated during infection with Ad5 mutants

We recently showed that infection with splicing-defective Ad5 mutants containing a deletion of the E4 region (ΔE4) leads to formation of viral dsRNA, as detected by immunofluorescence staining and RIP-seq using the dsRNA-specific J2 monoclonal antibody (Price *et al*, 2022a). This is accompanied by phosphorylation of dsRNA-activated kinase PKR during ΔE4 infection (Price *et al*, 2022a; Spurgeon & Ornelles, 2009). Additionally, we showed that infection with Ad5 mutants lacking the two VA RNAs (ΔVA) does not lead to detectable dsRNA formation with J2 antibody staining, despite prior literature which established that infection by this mutant results in PKR activation (Schneider *et al*, 1985; Kitajewski *et al*, 1986). To validate these observations further, we performed immunofluorescence using the dsRNA-specific monoclonal antibody 9D5, which is reportedly more sensitive than the widely used antibody J2 (Son *et al*, 2015). We evaluated dsRNA formation in A549 cells infected with wild-type (WT) Ad5, or the ΔVA and ΔE4 mutants (**Fig 1A**). Staining revealed faint cytoplasmic puncta representing mitochondrial dsRNA in uninfected cells, whereas large cytoplasmic aggregates were detected in cells transfected with the dsRNA analog poly(I:C) (Dhir *et al*, 2018; Corbet *et al*, 2022). In agreement with our recently reported results (Price *et al*, 2022a), robust nuclear staining was visualized in cells infected with the ΔE4 mutant, while no positive signal was detected for infections with Ad5 WT or ΔVA mutant (**Fig 1A**). Infection with mutants was further validated for the expression of VA RNAs I and II by primer extension analysis and expression of viral early and late proteins by immunoblotting (**Fig 1B,C**). Additionally, activation of the PKR pathway was confirmed by phosphorylation of PKR and eIF2α, using poly(I:C) transfection as a positive control (**Fig 1C**). These data demonstrate that although infection by both ΔVA and ΔE4 mutants induces PKR activation, formation of detectable dsRNA occurs only during ΔE4 infection.

**Figure 1:**
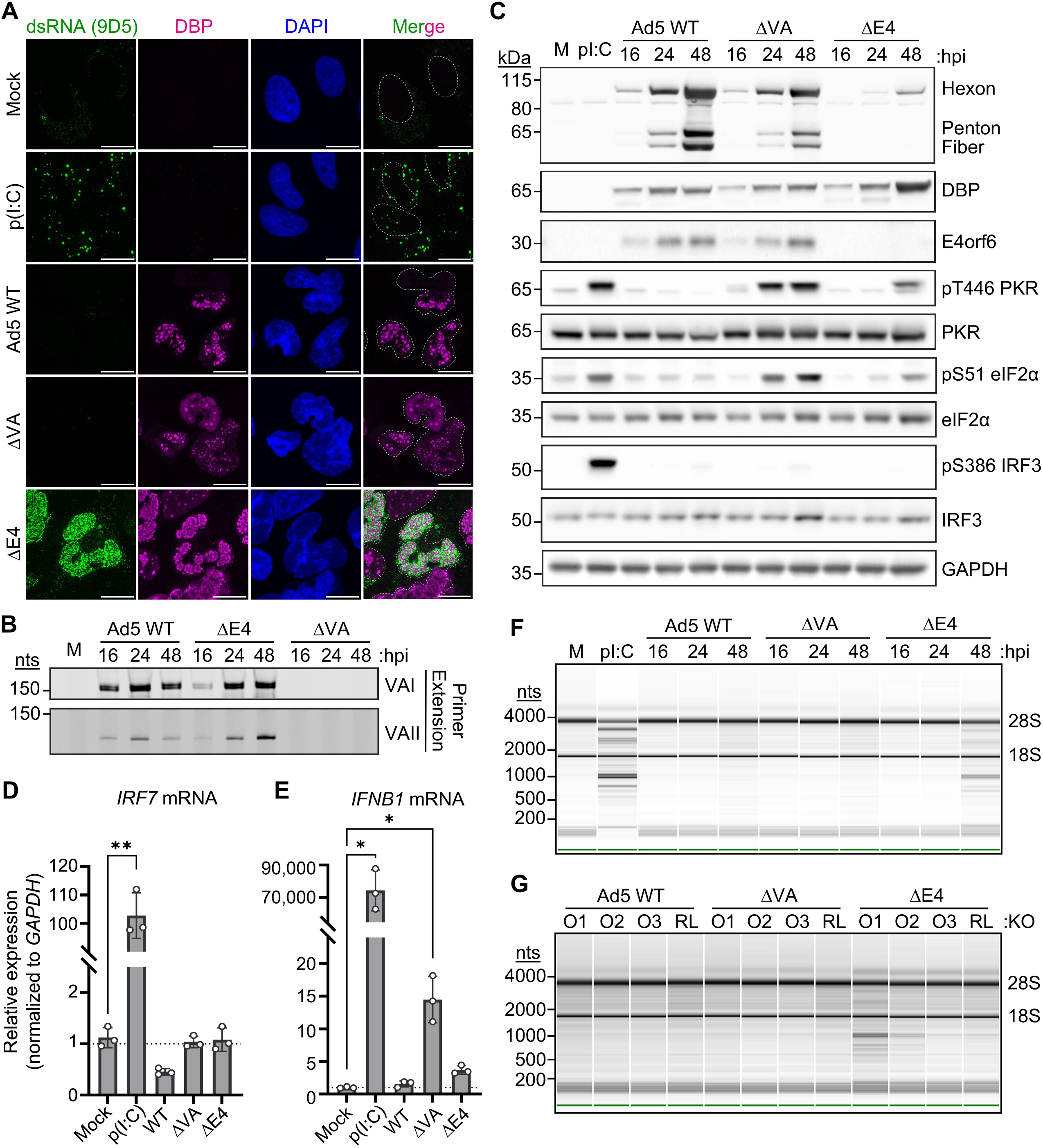
Activation of different dsRNA sensors during infection with Ad5 mutants. (**A**) A549 cells were transfected with poly(I:C) at 1 µg/mL for 6 h, mock-infected, or infected with Ad5 WT or ΔVA and ΔE4 mutants at an MOI of 10 for 48 h. Fixed cells were probed with the dsRNA-specific monoclonal antibody 9D5 (green), showing bright cytoplasmic staining upon poly(I:C) transfection and nuclear staining during infection with the ΔE4 mutant. The viral protein DBP (magenta) marks viral replication centers (VRCs). Nuclei were stained with DAPI (blue) and outlined by dashed white lines in merged images. Scale bar = 15 µm. (**B**) Whole cell RNA was prepared over a time course of infection and analyzed by PAGE. VA RNA expression was detected by primer extension analysis with fluorescently labeled, VA-specific primers. (**C**) Time-course immunoblot analysis of A549 cells infected with Ad5 WT or ΔVA and ΔE4 mutants. Poly(I:C) transfection was used as a positive control. PKR pathway activation was measured by autophosphorylation of PKR and phosphorylation of its downstream target eIF2α. RLR pathway activation was assessed by phosphorylation of IRF3. Viral proteins were detected with antibodies to early (DBP and E4orf6) and late proteins (Hexon, Penton, and Fiber). GAPDH was used as loading control. (**D-E**) Alternatively, RLR activation was measured by RT-qPCR for (**D**) *IRF7* and (**E**) *IFNB1* transcripts. A549 cells were infected or treated as in (**C**) and total RNA was extracted at 40 hpi and reverse transcribed. GAPDH-normalized data from three independent biological replicates are shown. Bars represent mean and error bars indicate standard deviation. Significance was calculated by one-way ANOVA followed by Dunnett’s multiple comparisons test, with * *P*<0.05, ** *P*<0.01 and non-significant results not displayed. (**F**) Total RNA was extracted from A549 cells treated as in (**C**) and run on BioAnalyzer to assess RNA integrity and RNase L activity. Results show ribosomal RNA cleavage following poly(I:C) transfection or infection with ΔE4, but not with WT or ΔVA mutant. (**G**) Alternatively, A549 lacking individual OAS isoforms (O1-3) or RNase L (RL) were infected with Ad5 WT or ΔVA and ΔE4 mutants. Total RNA was collected at 40 hpi and run on BioAnalyzer. Ribosomal RNA degradation indicates ΔE4-induced RNA cleavage is dependent on OAS3 and RNase L.

We next asked whether other dsRNA sensor pathways are activated during infection with WT or mutant viruses. We assessed RLR pathway activation by immunoblot analysis for IRF3 phosphorylated at serine 386, and RT-qPCR for measurement of *IRF7* and *IFNB1* transcripts. As expected, poly(I:C) transfection robustly induced IRF3 phosphorylation, consistent with previous reports (Gitlin *et al*, 2006; Kato *et al*, 2006, 2011). In comparison, phosphorylated IRF3 was barely detectable in cells infected with either Ad5 WT or mutant viruses (**Fig 1C**). Similarly, while poly(I:C) transfection strongly increased expression of *IRF7* and *IFNB1*, infection with the ΔVA mutant led to only modest increase in the expression of *IFNB1*, consistent with PKR activation (**Fig 1D,E**) (McAllister *et al*, 2012). The lack of induction of either IRF3 or IRF7 suggests that the RLR pathway is not activated during infection with Ad5 WT or the ΔVA and ΔE4 mutants.

Activation of the OAS1-3/RNase L pathway leads to characteristic degradation of 18S and 28S rRNAs, which can be visualized by bioanalyzer (Hovanessian *et al*, 1979; Silverman *et al*, 1983; Li *et al*, 2016). To determine whether this degradation occurs during infection with Ad5 mutants, total cellular RNA was extracted from infected cells at different timepoints or following poly(I:C) transfection, and samples were analyzed for rRNA integrity (**Fig 1F**). In line with prior observations, poly(I:C) transfection induced evident ribosomal RNA degradation (Li *et al*, 2016). RNA degradation was observed in late-stage ΔE4-infected samples but was absent during WT and ΔVA infections. Additionally, RNA degradation was completely abrogated during ΔE4 infection in A549 cells lacking RNase L or OAS3, but not in OAS1 or OAS2 knockout (KO) cells (**Fig 1G**). This is consistent with prior findings showing that OAS3 is the primary upstream dsRNA receptor during diverse viral infections (Li *et al*, 2016), despite recent studies suggesting that OAS1 and OAS2 may also promote context-dependent RNase L-mediated antiviral responses against certain viruses (Soveg *et al*, 2021; Merold *et al*, 2025).Collectively, these results suggest that formation of nuclear dsRNA during infection with ΔE4 mutant activates both PKR and RNase L pathways, while infection with ΔVA mutant does not yield detectable dsRNA and only activates the PKR signaling pathway.

### Infection with ΔVA and ΔE4 mutants induces formation of distinct cytoplasmic RNP granules

Upon activation, PKR promotes the formation of stress granules (SGs), while RNase L activation triggers the assembly of RNase L-dependent bodies (RLBs) (Watkins & Burke, 2024a). To investigate whether infection with Ad5 mutants induces granule formation following activation of dsRNA sensors, we used immunofluorescence to evaluate protein localization in A549 cells stained for G3BP1 and PABPC1 (**Fig 2A**). As a control for SG assembly, cells were treated with sodium arsenite (SA), which activates the eIF2α kinase HRI in response to oxidative stress (Pakos-Zebrucka *et al*, 2016). Alternatively, poly(I:C) was used for RNase L activation and induction of RLBs (Burke *et al*, 2020). No granules were detected in cells infected with Ad5 WT, whereas ΔVA infection induced the assembly of large cytoplasmic granules similar to SGs induced by SA treatment. In contrast, infection with the ΔE4 mutant led to the formation of small, spherical granules resembling RLBs assembled upon p(I:C) transfection (**Fig 2A**). In ΔE4 mutant infected cells, granule assembly was accompanied by nuclear accumulation of PABPC1, a hallmark of RNase L-mediated cytoplasmic mRNA decay (Burke *et al*, 2019, 2020). These observations were consistently reproduced across multiple cell lines (**Fig EV1**), indicating that infection with the ΔVA mutant produces SG-like granules, whereas the ΔE4 mutant promotes the assembly of RLB-like structures, with no granules being assembled during WT infection.

**Figure 2:**
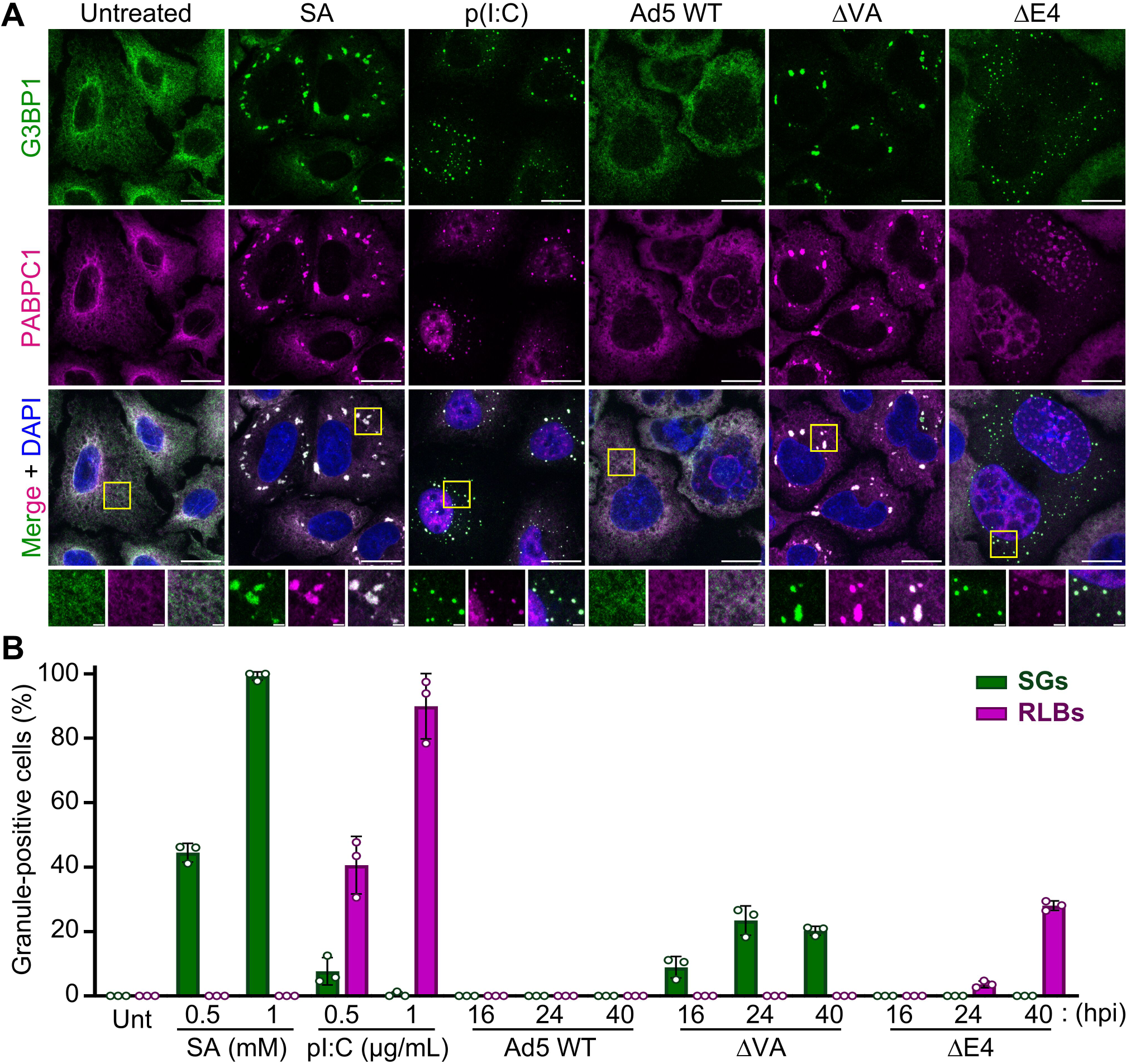
Infection with Ad5 mutants leads to assembly of granules similar to stress granules (SGs) and RNase L-dependent bodies (RLBs). (**A-B**) A549 cells were treated with sodium arsenite (SA, 0.5 or 1.0 mM for 1 h), poly(I:C) (0.5 or 1 µg/mL for 6 h) or infected with Ad5 WT, **Δ**VA or **Δ**E4 at an MOI of 10, as indicated. (**A**) Cells were fixed at 40 hpi and stained for granule markers G3BP1 (green) and PABPC1 (magenta), with nuclei stained in blue (DAPI). No granules were observed in untreated cells, while SA and poly(I:C) treatments induced the assembly of SGs and RLBs, respectively. Granule assembly was not detected upon WT infection, but SGs and RLBs were observed in ΔVA and ΔE4 infected cells, respectively. Scale bar = 15 µm (main panels), 2 µm (cropped panels). (**B**) Histogram showing the percentage of granule-positive cells over time following infection or increasing concentrations of SA or poly(I:C). Data are representative of three independent biological replicates. Bars represent mean, and error bars indicate standard deviation.

The formation of cytoplasmic RNP granules was further characterized and quantified over a time course of infection with Ad5 WT and mutants. The percentage of G3BP1 granule-positive cells was quantified by immunofluorescence in A549 cells, with SA or poly(I:C) treatments serving as controls for SG and RLB induction respectively (**Fig 2B**). Infection with the ΔVA mutant induced SG formation as early as 16 hours post infection (hpi), peaking at 24 hpi. Alternatively, assembly of RLBs during ΔE4 infection was only observed in a small proportion of cells by 24 hpi, increasing in frequency by 40 hpi. These kinetics closely correlate with the timing of PKR and RNase L activation in cells infected with these mutants (see **Fig 1C,F**). We noted that a considerable number of cells lacking obvious cytoplasmic G3BP1 foci also displayed nuclear PABPC1 signal during the late stage of WT and ΔVA infections (**Fig EV2A,B**) which resembled sites of nuclear speckle reorganization reported in Ad5 infected cells (Bridge *et al*, 1993, 1996; Gama-Carvalho *et al*, 2003). To address whether changes in PABPC1 localization impacted SG assembly, we also quantified the proportion of ΔVA-infected cells showing nuclear PABPC1, comparing cells with or without granules. Similar numbers of granules were observed in both groups, suggesting that nuclear PABPC1 localization does not impact granule assembly during ΔVA infection (**Fig EV2C**). Altogether, these findings demonstrate that ΔVA and ΔE4 mutants differentially induce the assembly of RNP granules with distinct kinetics, consistent with the differential activation of PKR and RNase L during infection.

### Virus-induced RNP granules reflect compositional differences between canonical SGs and RLBs

Despite being visually distinct, morphological differences alone are insufficient to distinguish SGs from RLBs, and viral infection may introduce additional context-dependent effects on granule composition. To address these limitations, we first sought to establish comparative reference proteomes for canonical SGs and RLBs under non-infectious conditions. Several studies have characterized SG components by proteomic approaches (Jain *et al*, 2016; Youn *et al*, 2018; Markmiller *et al*, 2018). In comparison, the RLB proteome has been examined in only one study, which used differential centrifugation of G3BP1-associated proteins in U2OS cells and suggested that, despite sharing many proteins with SGs, these granules also contain specific components (Burke *et al*, 2020). To explore further differences in protein composition between both condensates, we employed proximity labeling to identify proteins differentially enriched in granules upon SA or poly(I:C) treatments. For this, we used HEK293T cells with APEX2-GFP fused in-frame with the endogenous *G3BP1* locus (**Fig EV3A,B**) (Elmsaouri *et al*, 2022; Markmiller *et al*, 2018). Proteomic datasets showed consistent normalization and reproducibility across conditions and replicates (**Fig EV3C-G**). Comparison of SG-enriched proteins identified in our analysis with a previous dataset published using the same system showed significant overlap of identified proteins, validating our experimental approach (**Fig EV3H**). In contrast, comparison with the previous RLB dataset showed limited overlap, which could reflect cell-type specific differences in protein recruitment or methodological biases between fractionation and proximity labeling techniques (**Fig EV3I**).

Analysis of the SG and RLB datasets for differential protein enrichment revealed substantial overlap but also clear differences in granule composition (**Fig 3A,B** and **Table EV1**), including proteins previously reported to be present in both granules (e.g. UBAP2L, Caprin1) or uniquely enriched in SGs (e.g. FAM120A, TIA1) (Burke *et al*, 2020; Burke, 2023). We validated our proteomics data by immunofluorescence to assess the recruitment of proteins to SGs and RLBs in cells treated with SA or poly(I:C), and examined these differences in granules induced by viral mutants (**Fig 3C**,**D** and **EV4A,B**). We confirmed the accumulation of FXR1, FMRP, Caprin1 and UBAP2L in both granules, whereas FAM120A, eIF4G1, PRRC2C and Ago2 localized exclusively to SGs induced by either SA or **Δ**VA infection. LSm14A was significantly more enriched in the RLB dataset, showing strong staining in granules induced by poly(I:C) and ΔE4 infection, but only partial relocalization from P-bodies to SGs. To determine whether this effect is specific to LSm14A, we also probed cells for the P-body marker Dcp1b. Lack of colocalization between Dcp1b and G3BP1-stained RLBs confirmed the specific ability of LSm14A to shuttle between P-bodies, SGs and RLBs (**Fig EV4C**). Collectively, these results reinforce previously documented differences between SGs and RLBs, and they suggest that granules induced during adenovirus mutant infection closely resemble those formed in response to chemical stressors, although additional studies will be required to establish their full proteomic equivalence in infected cells.

**Figure 3:**
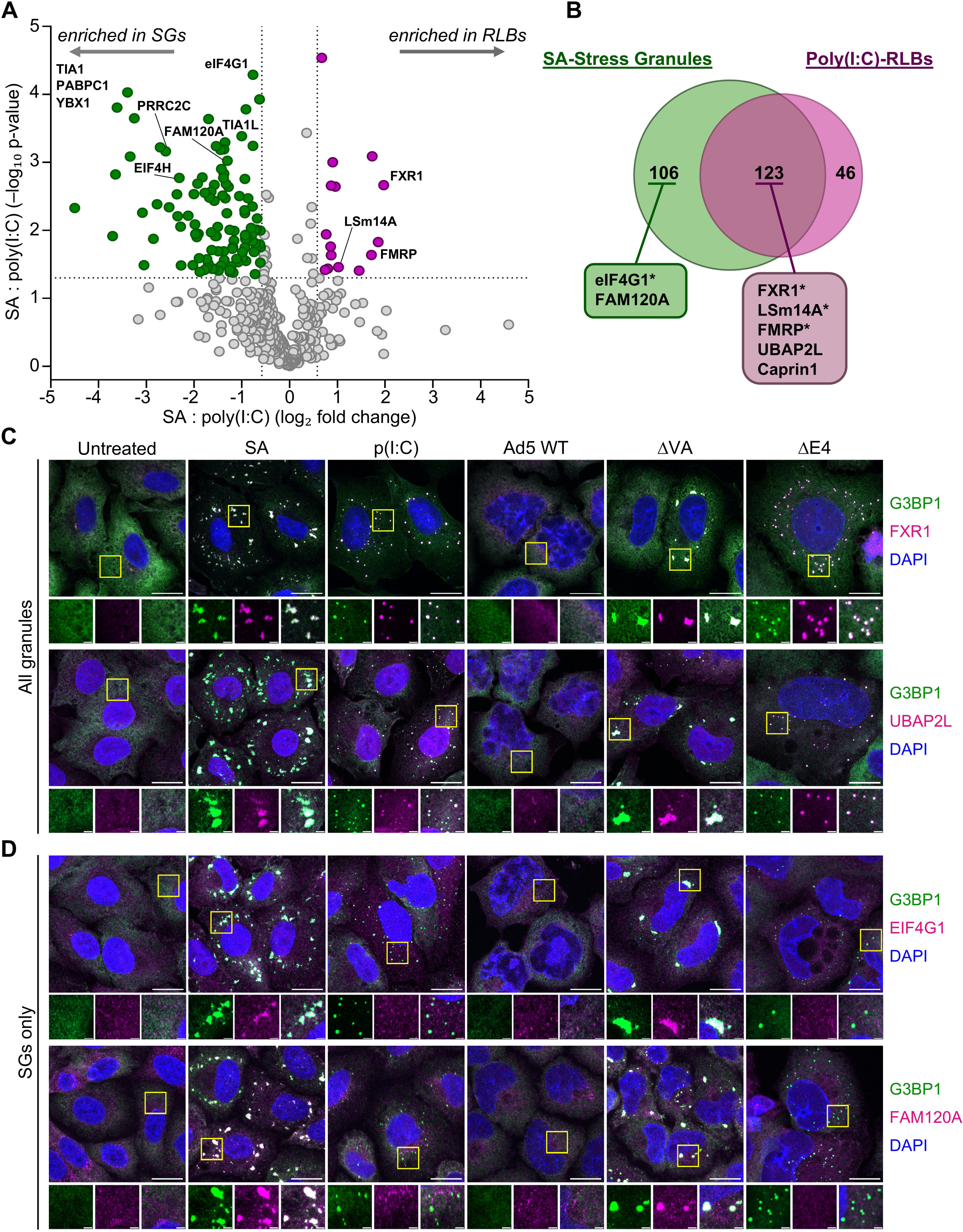
Proteomic analysis reveals differential enrichment of proteins in SGs and RLBs. (A) Volcano plot showing differentially enriched proteins following APEX proximity labeling in HEK293T cells expressing G3BP1-APEX2-GFP fusion protein. Cells were treated with sodium arsenite (SA, 1.0 mM for 1 h) or poly(I:C) (1.0 µg/mL for 4 h) to induce granule formation before labeling. The x-axis shows the log_2_ fold-change of the APEX abundances, normalized to input, for SA compared to poly(I:C). The y-axis indicates the –log_10_ p-value for the comparison of normalized APEX abundances for the three biological replicates of SA and poly(I:C) treatments. Proteins significantly enriched in SGs (left, green) or RLBs (right, magenta) are highlighted. Proteins validated by our proteomic analysis or identified in previously published datasets are labeled in the figure. (**B**) Venn diagram comparing proteins enriched in SGs and RLBs. Proteins that are unique to SGs (left, green), unique to RLBs (right, magenta), or shared between the granules (intersection) are indicated. Proteins further characterized in this study are highlighted by asterisks. (**C-D**) Validation of APEX-identified proteins was performed in A549 cells treated with sodium arsenite (SA, 0.5 mM for 1 h), poly(I:C) (0.5 µg/mL for 6 h) or infected with Ad5 WT, **Δ**VA or **Δ**E4 viruses at an MOI of 10 for 40 h. G3BP1 staining (green) was used as a marker for granule assembly. (**C**) Co-staining with FXR1 or UBAP2L (magenta) confirms recruitment of proteins to both SGs and RLBs. (**D**) Staining for eIF4G1 or FAM120A (magenta) shows proteins recruited to SGs, but absent from RLBs. Nuclei were stained with DAPI (blue). Scale bar = 15 µm (main panels), 2 µm (cropped panels).

Unlike stress granules, RLBs have been shown to assemble independently of G3BP1 and G3BP2 proteins (Burke *et al*, 2020). To evaluate whether granules induced during infection share this property, we infected A549 G3BP1/2 KO cells with Ad5 WT or mutants, using SA and poly(I:C) treatments as controls. Cells were stained for FXR1 and PABPC1, confirming the assembly of RLBs in response to either poly(I:C) or ΔE4 infection (**Fig EV5A**). In comparison, stress granules did not form in response to SA or ΔVA infection, consistent with G3BP proteins being required for SG maturation. Previous work has also shown that UBAP2L contributes to the scaffolding of stress granules (Asano-Inami *et al*, 2023; Yang *et al*, 2020). In cells lacking G3BP proteins, UBAP2L forms small nucleating granules that fail to mature into full SGs (Cirillo *et al*, 2020). To distinguish these structures from RLBs in the G3BP1/2 KO background, we co-stained cells for UBAP2L and FXR1. In SA-treated cells, UBAP2L granules showed minimal FXR1 recruitment, whereas RLBs displayed robust FXR1 staining during both poly(I:C) transfection and ΔE4 infection (**Fig EV5B**). Together, these findings demonstrate that ΔE4-induced granules are similar to poly(I:C)-induced RLBs, recruiting similar protein markers and assembling in cells lacking G3BP proteins.

### PKR and RNase L show distinct requirements for formation of virus-induced RNP granules

To establish whether virus-induced granules form downstream of cellular dsRNA sensor pathways, we evaluated granule assembly upon infection in A549 cells individually knocked out for PKR or RNase L. Immunoblot analysis confirmed the loss of PKR and RNase L expression and successful infection of each cell line with Ad5 WT and mutant viruses (**Fig 4A**). We then assessed granule formation in response to chemical stressors or viral infections by staining cells for G3BP1 and PABPC1, followed by quantification of granule-positive cells (**Fig 4B,C**). No granules were observed in untreated or WT infected cells. As expected, SG formation during SA treatment was unaffected by the absence of PKR or RNase L expression. By comparison, RLB assembly occurred independently of PKR expression following poly(I:C) transfection in PKR KO cells. In RNase L KO cells, however, poly(I:C) induced the formation of large SGs, consistent with previous findings indicating that poly(I:C) triggers SG assembly in RNase L-deficient cells in response to PKR activation (Burke *et al*, 2020). During ΔVA infection, granules were visualized in parental and RNase L KO cells, but not in PKR KO cells. This suggests SGs form downstream of PKR activation in cells infected by the ΔVA mutant. Unexpectedly, ΔE4-induced cytoplasmic granules were detected in all cell lines, including those lacking RNase L (**Fig 4B,C**).

**Figure 4:**
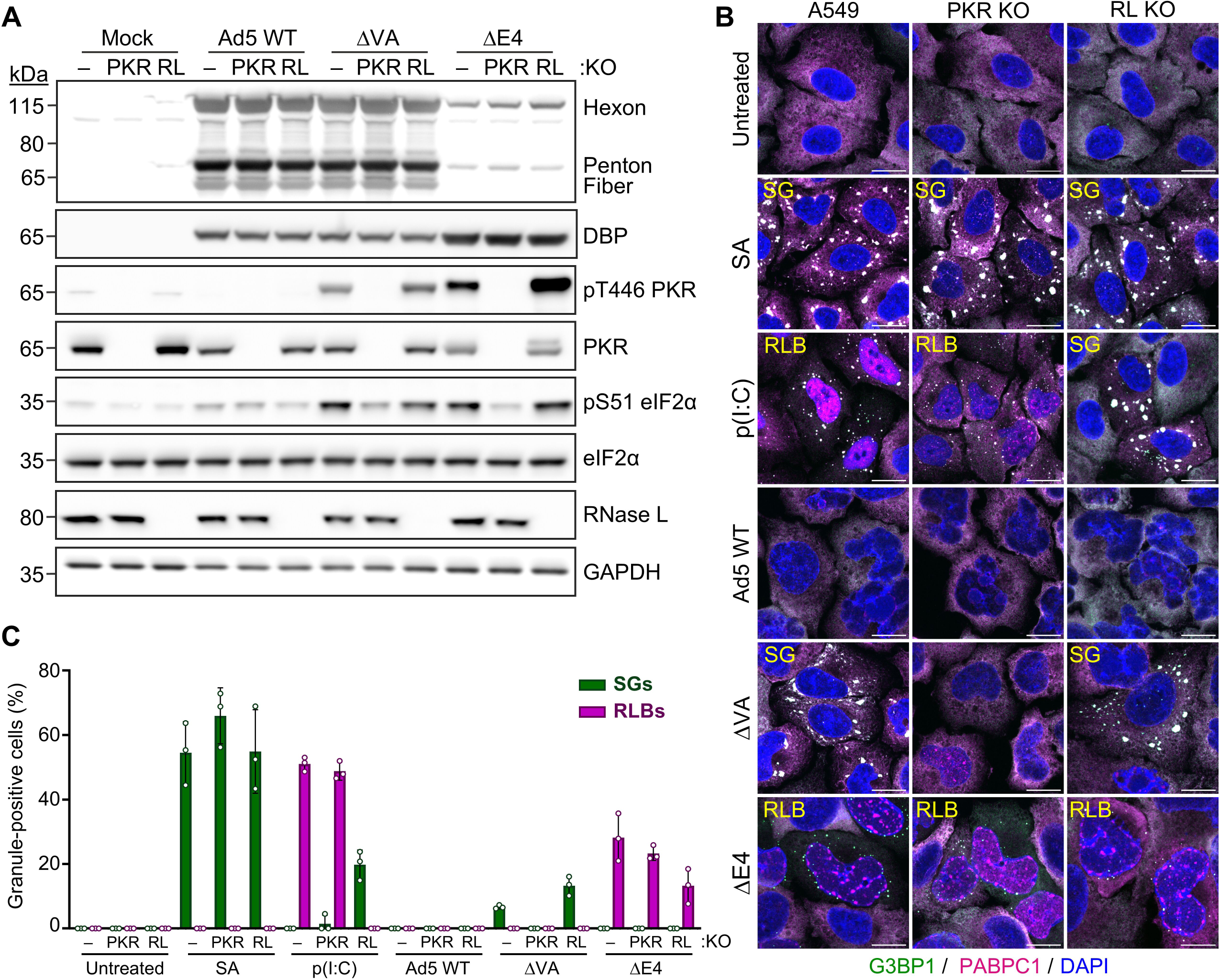
Requirements for granule formation during mutant AdV infection. (**A**) Immunoblot analysis of viral late proteins, RNase L expression, and PKR pathway activation in parental (–) and KO A549 cell lines lacking PKR and RNase L (RL). GAPDH was used as a loading control. Cells were infected with Ad5 WT, **Δ**VA and **Δ**E4 at an MOI 10 for 40 h. (**B**) Panels show merged IF images for granule markers G3BP1 (green) and PABPC1 (magenta) in the indicated A549 cell lines (parental, PKR KO, RNase L KO) following chemical stress with sodium arsenite (SA, 0.5 μM 1 h) and poly(I:C) (0.5 μg/mL 6 h) or viral infection for 40 h. Nuclei are stained with DAPI (blue). Scale bar = 15 µm. The proportion of cells displaying granules is quantified in (**C**). Results are representative of three independent biological replicates. Bars represent mean, and error bars indicate standard deviation.

Furthermore, granule assembly was accompanied by nuclear relocalization of PABPC1, as observed in parental cells (**Fig 4B,C**). Notably, this occurred despite the lack of RNase L activity, as confirmed by the absence of ribosomal RNA degradation in these cells (see **Fig 1G**). Similar results were observed in U2OS RNase L KO cells following poly(I:C) transfection or ΔE4 infection (**Fig EV6A,B**). These findings demonstrate that ΔVA infection induces PKR-dependent SGs, whereas ΔE4 infection promotes the assembly of RLB-like granules which are independent of RNase L activity.

An additional hallmark of RNase L activity is the widespread cleavage of cytoplasmic RNAs, which drives RLB assembly and bulk PABPC1 translocation to the nucleus (Burke *et al*, 2022). To determine whether granule assembly during ΔE4 infection is accompanied by cytoplasmic mRNA turnover, we performed single-molecule fluorescent *in situ* hybridization (smFISH) for *GAPDH* transcripts in parental and RNase L KO cells infected with Ad5 WT or the ΔE4 mutant, or transfected with poly(I:C) (**Fig 5A-D**). Consistent with previous findings, *GAPDH* mRNA signal was strongly reduced in response to poly(I:C) (Burke *et al*, 2019). Reduced *GAPDH* mRNA signal was also observed in ΔE4-infected cells. In contrast, *GAPDH* signal during WT infection was equivalent to signal quantified in uninfected cells (**Fig 5A,B**), suggesting that the reduction in *GAPDH* mRNA in ΔE4-infected cells is a result of RNase L activation. Supportive of this interpretation, *GAPDH* mRNA levels were comparable across all conditions in RNase L KO cells, with a slight increase in signal during poly(I:C) transfection and ΔE4 infection (**Fig 5C,D**). This confirms that the global degradation of cytoplasmic mRNAs requires RNase L activity in infected cells. Collectively, these data indicate that ΔE4-induced granules accompanied by nuclear PABPC1 translocation do not depend on widespread mRNA degradation, since there was no evidence of *GAPDH* mRNA turnover in infected RNase L KO cells. This suggests the existence of additional mechanisms for granule assembly during infection with the ΔE4 mutant virus.

**Figure 5:**
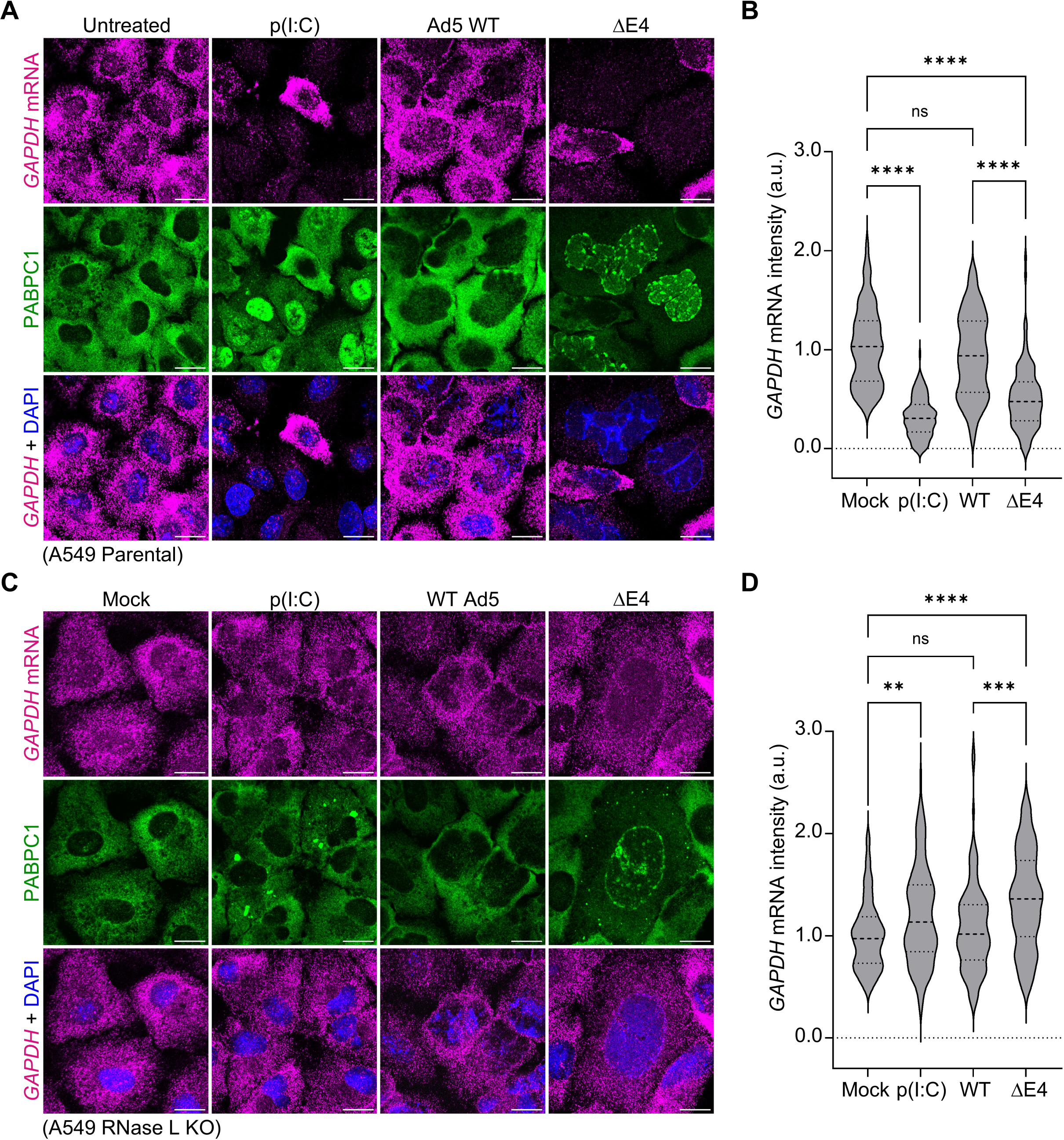
Widespread mRNA turnover depends on RNase L in ΔE4-infected cells. **(A-D)** A549 parental or RNase L KO cells were treated with poly(I:C) (1 µg/mL for 6 h) or infected with Ad5 WT or **Δ**E4 (MOI 10 for 40 h). Cells were stained by smFISH for *GAPDH* transcripts (magenta), PABPC1 (green) to indicate granule assembly and nuclei is stained with DAPI (blue). Scale bar = 15 µm. *GAPDH* smFISH intensity was quantified per cell as corrected total cell fluorescence (CTCF = Area × [Mean intensity − Mean background]). Violin plots show signal distribution per cell with median and quartiles indicated by dashed lines. (**A, B**) In parental cells, loss of *GAPDH* signal was observed upon poly(I:C) transfection and **Δ**E4 infection. Quantification was performed in mock n=133, poly(I:C) n=116, Ad5 WT n=169 and **Δ**E4 n=106. (**C, D**) In RNase L KO cells, *GAPDH* mRNA signal was observed among all conditions. The total number of cells analyzed was mock n=142, poly(I:C) n=118, Ad5 WT n=103 and **Δ**E4 n=77. For poly(I:C) and **Δ**E4 conditions, only cells displaying granules or nuclear PABPC1 localization were quantified. Statistical significance was measured by Kruskal-Wallis test followed by Dunn’s multiple comparisons test, with ns = not significant, ** = *P*<0.01, *** = *P*<0.001, **** = *P*<0.0001.

### Cytoplasmic granules induced by ΔE4 infection share properties with canonical RLBs

To characterize further the extent to which ΔE4-induced granules are similar to canonical RLBs, we performed immunofluorescence in RNase L KO cells following poly(I:C) transfection or ΔE4 infection. In these cells, granules assembled in response to ΔE4 infection showed recruitment of UBAP2L and FXR1 in a similar way to poly(I:C)-induced granules (**Fig 6A**). In contrast, co-staining of G3BP1 with eIF4G1 or Dcp1b confirmed ΔE4-induced granules are distinct from SGs and P-bodies (**Fig 6B,C**). Consistent with prior reports showing that RLBs contain polyadenylated RNA, we also showed by poly(A)+ FISH that ΔE4-induced granules stain for this marker, despite lack of widespread mRNA turnover (**Fig 5C,D** and **6D**) (Burke *et al*, 2020). Another difference between RLBs and SGs is their requirement for active translation during granule assembly. SG formation requires polysome disassembly, which is blocked by the translation inhibitor cycloheximide (CHX) (Mollet *et al*, 2008). In comparison, CHX treatment does not block RLB assembly as RNase L also cleaves mRNAs bound to polysomes (Burke *et al*, 2019). To assess the impact of CHX on RNP granule assembly, parental and RNase L KO cells were transfected with poly(I:C) or infected with ΔE4 and stained for granule markers. Poly(I:C) induction of RLBs in parental cells was not affected by translation arrest, while the assembly of SGs was completely blocked in RNase L KO cells. By contrast, ΔE4 infection induced granules resembling RLBs in all conditions, which was accompanied by nuclear staining for PABPC1 (**Fig 6E,F**). Together, these findings indicate that granules induced during ΔE4 infection in RNase L KO cells are distinct from SGs, while still sharing compositional and biochemical hallmarks of canonical RLBs.

**Figure 6:**
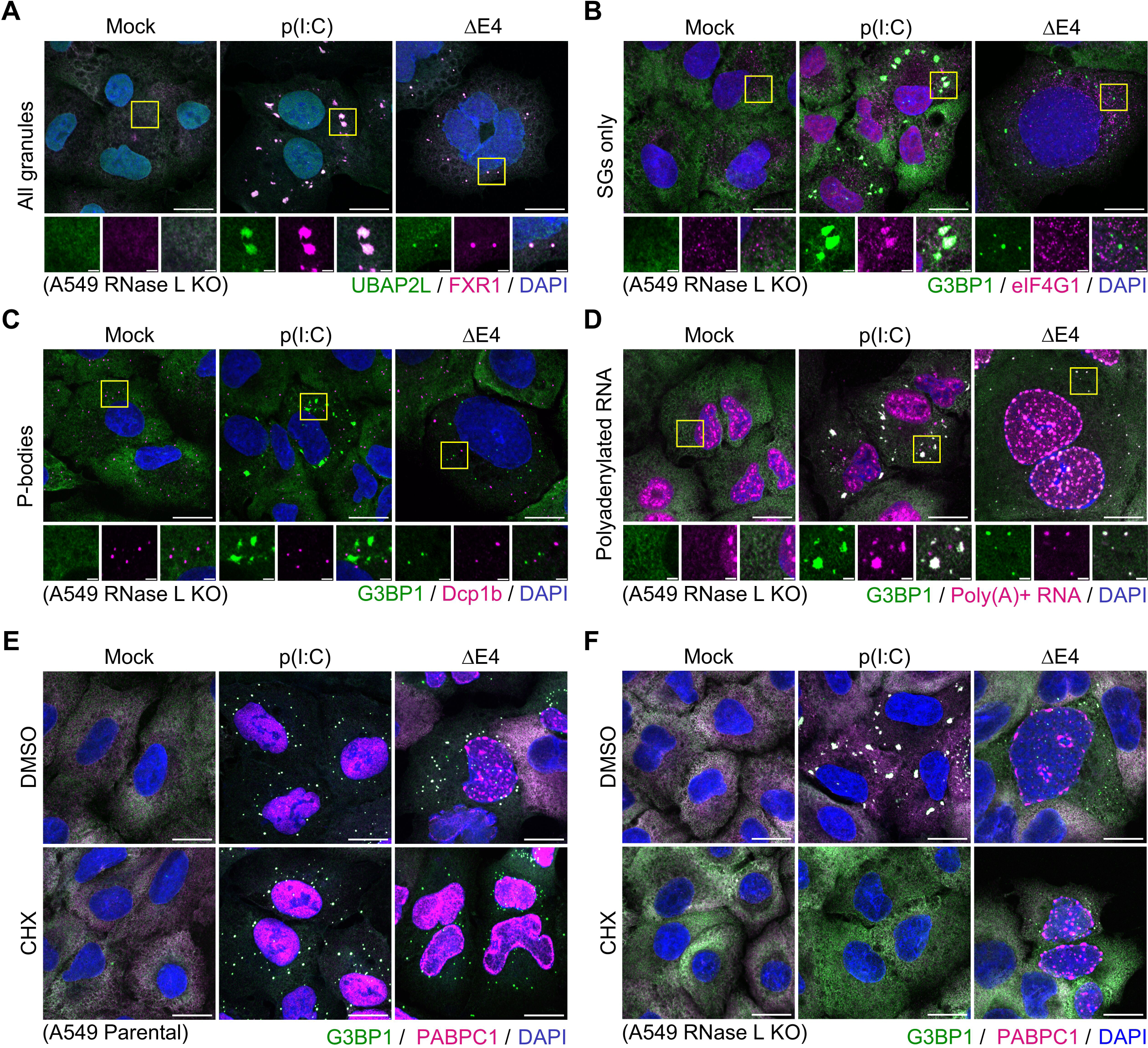
Granules induced by ΔE4 infection in RNase L KO cells share structural and biochemical properties with RLBs. (**A-D**) A549 RNase L KO cells treated with poly(I:C) (1 µg/mL for 6 h) or infected with ΔE4 (MOI 10 for 40 h) were stained for different granule markers. (**A**) UBAP2L (green) and FXR1 (magenta) show colocalization of both proteins to SGs induced by poly(I:C) and RLB-like granules induced during **Δ**E4 infection. Alternatively, the co-staining of G3BP1 with eIF4G1 (**B**) or Dcp1b (**C**) confirms **Δ**E4-induced RLBs do not colocalize with SG-unique proteins or P-bodies. (**D**) Staining for G3BP1 (green) and detection of polyadenylated RNA by FISH (magenta) shows recruitment of poly(A)+ RNA to both SGs and **Δ**E4-induced granules. (**E**) A549 parental and RNase L KO cells (**F**) were transfected with poly(I:C) (1 µg/mL for 6 h) or infected with **Δ**E4 (MOI 10 for 40 h). Cells were treated with DMSO or cycloheximide (CHX) 100 µg/mL to block translation and stress granule assembly. Fixed cells were stained for G3BP1 (green) and PABPC1 (magenta), with DAPI staining nuclei (blue). CHX treatment does not block poly(I:C) induced RLBs in parental cells, but blocks SG assembly in RNase L KO cells. **Δ**E4 infection induced RLB-like granules independently of CHX treatment in both cell lines. Scale bar = 15 µm (upper panels), 2 µm (cropped panels).

### Infection with ΔE4 mutant leads to translation arrest independent of PKR and RNase L

In addition to RNP granule induction, activation of PKR and RNase L leads to global translational arrest dependent on eIF2α phosphorylation or RNA cleavage, respectively (Burke *et al*, 2019; Pakos-Zebrucka *et al*, 2016). To probe further this phenotype, A549 parental or RNase L KO cells were transfected with poly(I:C) or infected with ΔE4 and briefly pulsed with puromycin prior to fixation. PABPC1 staining was used to mark granule assembly, while puromycin incorporation was used to monitor active translation (**Fig EV7A,B**). As expected, robust puromycin signal was observed in mock or granule-negative cells, whereas puromycin incorporation was markedly reduced in poly(I:C) and ΔE4-infected cells displaying granules. Since ΔE4 infection in RNase L KO cells also activates PKR, we asked whether PKR activity is dispensable for induction of ΔE4-dependent granules and translation shutoff in cells lacking RNase L expression. Using A549 PKR/RNase L double-KO cells, we verified that ΔE4 infection could still induce the assembly of cytoplasmic granules similar to RLBs, which was accompanied by nuclear PABPC1 staining (**Fig 7A**). In contrast, poly(I:C) failed to induce either SGs or RLBs in cells lacking both dsRNA sensors. Puromycin pulsing in double-KO cells revealed that formation of **Δ**E4-induced RNP granules is accompanied by a strong reduction in puromycin incorporation in comparison to uninfected or poly(I:C)-transfected cells (**Fig 7B**). Importantly, this effect is observed despite lack of detectable eIF2α phosphorylation in these cells (**Fig 7C**). To rule out any effect associated with eIF2α, double-KO cells treated with SA, poly(I:C) or infected with the **Δ**E4 mutant were treated with ISRIB, which restricts the inhibitory effect of phosphorylated eIF2α on translation initiation (Sidrauski *et al*, 2015). Consistent with previous reports, ISRIB completely prevented the assembly of SGs during SA treatment, restoring translation in these cells (**Fig 7D,E**). In comparison, no rescue of puromycin incorporation was observed in **Δ**E4-infected cells displaying RNP granules, indicating that translation arrest in these cells does not depend on eIF2α phosphorylation. Collectively, these findings demonstrate that ΔE4 infection induces translational repression and assembly of granules independently of PKR, RNase L and eIF2α phosphorylation. This suppports the existence of a non-canonical pathway linking the accumulation of viral nuclear dsRNA to cytoplasmic RNP granule assembly and translational control.

**Figure 7:**
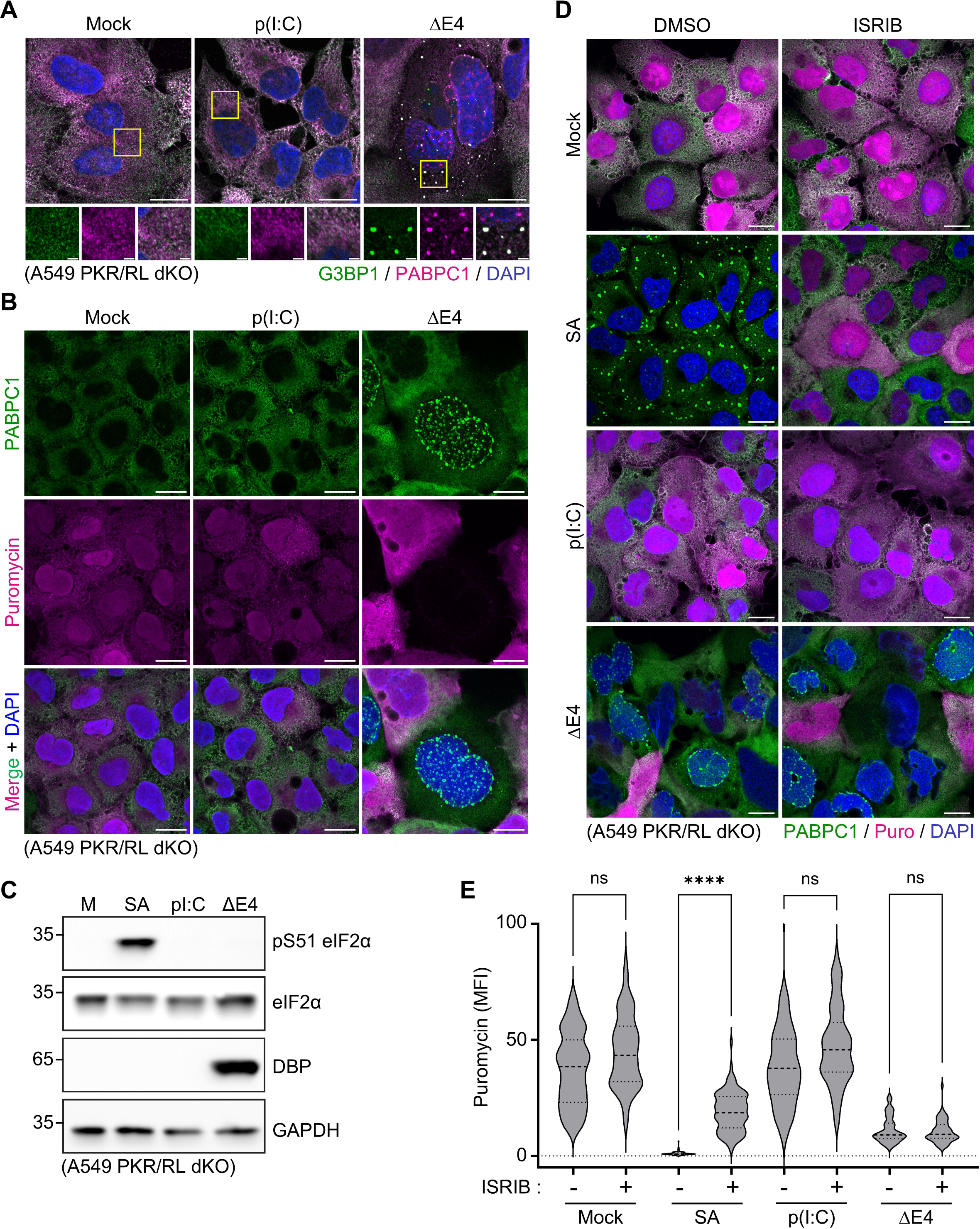
Granule assembly and translational arrest induced by ΔE4 infection is not dependent on RNase L and PKR. A549 PKR / RNase L double-KO cells were treated with poly(I:C) (1 µg/mL for 6 h) or infected with **Δ**E4 (MOI 10 for 40 h). (**A**) Cells were stained for G3BP1 (green) and PABPC1 (magenta). **Δ**E4-induced granules are visualized in the cytoplasm, with PABPC1 nuclear staining. No granules were assembled upon poly(I:C) transfection. (**B**) To assess active translation, cells were treated with puromycin (10 µg/mL) for 10 min before fixation. PABPC1 staining (green) is shown to indicate granule assembly. Puromycin incorporation (magenta) is visualized in mock or poly(I:C)-treated cells, but not in **Δ**E4-infected cells displaying RNP granules. (**C-D**) A549 PKR / RNase L double-KO cells were treated with SA (0.5 mM for 1 h), poly(I:C) (1 µg/mL for 6 h) or infected with **Δ**E4 (MOI 10 for 40 h). (**C**) Immunoblotting analysis showing eIF2α phosphorylation upon SA treatment, but not in response to poly(I:C) and **Δ**E4 infection. (**D**) Alternatively, cells were further treated with DMSO or ISRIB (200 nM) and incubated with puromycin (10 µg/mL) for 10 min before fixation. ISRIB blocked SA-induced SG assembly and partially restored translation upon stress, but showed no effect in **Δ**E4-infected cells, as quantified in (**E**). Violin plots show puromycin incorporation per cell quantified by mean fluorescence intensity (MFI), with median and quartiles shown as dashed lines. For DMSO, we analyzed mock n=80, SA n=89, poly(I:C) n=66, and **Δ**E4 n=59. For ISRIB treatment, we analyzed mock n=92, SA n=94, poly(I:C) n=72, and **Δ**E4 n=60. For **Δ**E4 infection, puromycin signal was quantified in cells displaying granules and nuclear PABPC1 localization. Statistical significance was measured by Kruskal-Wallis test followed by Dunn’s multiple comparisons test, with ns = not significant, **** = *P*<0.0001. Nuclei were stained with DAPI (blue). Scale bar = 15 µm (main panels), 2 µm (cropped panels).

## DISCUSSION

Adenovirus infection has been employed for many years to investigate the interplay between dsRNA formation and RNA sensing mediated by PKR (Maran & Mathews, 1988; Jacobs & Langland, 1996). Activation of PKR during infection with viral mutants lacking VA RNAs (ΔVA) has been speculated to result as a consequence of bidirectional transcription from both viral genomic DNA strands (Siekierka *et al*, 1985; Mellits *et al*, 1990). We recently showed that dsRNA is formed in the nucleus upon AdV infection with splicing-defective E4-deleted mutants (ΔE4), but is not observed during infection with Ad5 WT or ΔVA mutants (Price *et al*, 2022a). Here, we show that these mutants differentially engage dsRNA sensing pathways leading to distinct downstream outcomes. We verified that infection with the ΔVA mutant strongly activated PKR, but not other dsRNA sensors. In contrast, ΔE4 mutant infection induced both PKR and OAS3/RNase L pathways, consistent with dsRNA formation by this viral mutant. Notably, none of our mutants led to activation of the RLR pathway, as evidenced by the absence of IRF3 phosphorylation or expression of other downstream genes. Lack of RLR responses, in parallel with the activation of other dsRNA sensors, could reflect differences intrinsic to each mutant, since ΔVA infection does not produce detectable dsRNA, and ΔE4-induced dsRNA is largely restricted to nuclear sites, limiting the access of cytoplasmic receptors RIG-I and MDA5 (Kato *et al*, 2011; Rehwinkel & Gack, 2020; Price *et al*, 2022a). However, we cannot exclude the influence of other viral elements in restricting this pathway, as previously described for E1A proteins (Leonard & Sen, 1996; Fonseca *et al*, 2012; Zemke & Berk, 2017; Olanubi *et al*, 2017). Collectively, these findings exemplify how diverse viral elements act in separate axes to evade cellular responses to dsRNA, also pointing to PKR and RNase L as the major players engaged during mutant AdV infection.

We discovered that activation of distinct dsRNA sensors by AdV mutants resulted in differential assembly of cytoplasmic RNP condensates. Infection with the ΔVA mutant triggered assembly of PKR-dependent cytoplasmic granules, mirroring SGs formed by other canonical stress responses mediated by eIF2α phosphorylation (Brownsword & Locker, 2023; Eiermann *et al*, 2020; Pakos-Zebrucka *et al*, 2016; McCormick & Khaperskyy, 2017). Conversely, ΔE4 mutant infection induced small, rounded cytoplasmic granules resembling RLBs, correlating with RNase L activation (Burke *et al*, 2019). In addition to their different morphologies, these RNP granules have been reported to have distinct protein compositions (Burke *et al*, 2020; Burke, 2023). Such variation is not restricted to the type of granule (i.e. SGs and RLBs), with changes reported in the proteome of stress granules depending on the cell type and nature of the stress stimulus, which include oxidative stress, UV irradiation and viral infection (Piotrowska *et al*, 2010; Aulas *et al*, 2017; Markmiller *et al*, 2018; Zhou *et al*, 2024). Here we further characterized the differences between SGs and RLBs by proteomic and immunofluorescence analysis. We identified several proteins shared between both condensates, while others were uniquely recruited to SGs, consistent with previous studies (Burke *et al*, 2020; Burke, 2023). Notably, these compositional differences were recapitulated for selected granule markers during viral infection, suggesting viral-induced granules are similar to their chemically-induced counterparts produced in response to SA or poly(I:C) transfection. Together, these findings demonstrate that the activation of different dsRNA sensors directs cellular stress responses into the formation of RNP granules with distinct compositions, which is mimicked during infection with AdV mutants.

Although similar properties were observed for RLBs induced by ΔE4 and poly(I:C), granules produced during ΔE4 infection were assembled independently of RNase L. In RNase L knockout cells, poly(I:C) induced the assembly of PKR-dependent SGs, whereas ΔE4 infection formed granules similar in shape and composition to canonical RLBs. The assembly of these RLB-like granules was not accompanied by widespread cleavage of ribosomal and cytoplasmic mRNA, confirming the absence of RNase L catalytic activity (Burke *et al*, 2019). We further explored these observations in PKR and RNase L double-KO cells, confirming that RNP granule assembly during ΔE4 infection does not depend on either of these dsRNA sensors. The assembly of RLB-like structures has also been reported during infection by several cytoplasmic RNA viruses, including flaviviruses such as dengue (DENV-2), West Nile (WNV), and Zika (ZIKV) viruses, as well as coronaviruses like SARS-CoV-2 (Burke *et al*, 2021; Watkins & Burke, 2024b; Burke, 2023). In contrast to the ΔE4 adenovirus mutant, flavivirus infection promotes SG assembly in RNase L-deficient cells, whereas SARS-CoV-2 fails to form SGs in this context, likely due to viral proteins that antagonize the scaffolding functions of G3BP1 and G3BP2 (Zheng *et al*, 2021; Liu *et al*, 2022). These contrasting outcomes illustrate how formation of dsRNA in specific cellular compartments during viral infection can differentially modulate the assembly of RNP granules. Our findings suggest that ΔE4-mediated granule assembly can occur independently of major cellular dsRNA sensors such as PKR and RNase L.

Translational arrest was also observed during ΔE4 infection in double-KO cells, indicating PKR and RNase L are not required for this process. Previous reports have demonstrated that several viruses promote translational shutoff by activating other eIF2α kinases, such as GCN2 or PERK (Zhang *et al*, 2019; Brownsword & Locker, 2023; Eiermann *et al*, 2020). To determine whether ΔE4-induced translational arrest was driven by a different eIF2α kinase, we confirmed that eIF2α is not phosphorylated during ΔE4 infection in PKR/RNase L double-KO cells. This observation was further supported by treating these cells with ISRIB, an antagonist of phosphorylated eIF2α (Sidrauski *et al*, 2015; Zyryanova *et al*, 2018; Rabouw *et al*, 2019). ISRIB treatment promoted the disassembly of stress granules and restored translation upon SA exposure, but showed no effect in ΔE4-infected cells. These results suggest that ΔE4 infection can also drive translational repression through additional non-canonical pathways. We note that production of nuclear dsRNA during ΔE4 mutant infection constitutes an unconventional site for dsRNA detection, which typically occurs in the cytoplasm during infection by most viruses or after poly(I:C) transfection (Burke, 2023; McCormick & Khaperskyy, 2017; Price *et al*, 2022a; Corbet *et al*, 2022, 2023). Therefore, it is tempting to consider that nuclear dsRNA activates alternative cellular sensors, leading to granule assembly and translation shutoff through non-canonical mechanisms. Consistent with this idea, the phenotype observed during ΔE4 infection in KO cells resembles the effects of nucleolytic activity mediated by RNase L, despite the lack of widespread RNA degradation. The hypothetical activation of a different endonuclease in response to nuclear dsRNA could explain the assembly of RLB-like granules accompanied by PABPC1 nuclear relocalization, as well as translation shutoff by eIF2α-independent mechanisms. Although no changes in *GAPDH* mRNA levels were detected upon ΔE4 infection in RNase L KO cells, we cannot exclude the possibility that specific transcripts or RNA isoforms are targeted for cleavage during infection, which could lead to the observed outcomes. This possibility is supported by previous reports describing that selective cleavage of tRNAs by angiogenin or SLFN proteins can elicit translational arrest, with the latter associated with cellular antiviral responses (Yamasaki *et al*, 2009; Emara *et al*, 2010; Yang *et al*, 2018; Kim & Weitzman, 2022). Such an effect would be masked by RNase L activity in parental cells, since infection with the ΔE4 mutant coincides with other hallmarks of RNase L activation (Donovan *et al*, 2017). However, additional studies are necessary to elucidate whether nuclear dsRNA alone is responsible for inducing these cellular responses.

In summary, our work describes the activation of distinct dsRNA sensing pathways during infection with mutant adenoviruses, leading to divergent cellular outcomes. Activation of PKR by the ΔVA mutant or both PKR and RNase L during ΔE4 infection illustrates how different viral elements counter host antiviral defenses. Activation of these pathways culminates in the formation of RNP granules with distinct compositions, which share biochemical and structural features with canonical SGs and RLBs. Notably, the ability of the ΔE4 mutant to promote granule assembly and translational repression independently of PKR, RNase L, or eIF2α phosphorylation indicates the existence of alternative cellular responses triggered during viral infection or nuclear dsRNA formation. Further characterization of these non-canonical pathways will be essential to understand the full landscape of how cells sense and respond to stressors. Together, our findings broaden the current view of cellular surveillance to dsRNA during viral infection, exploring the interplay between well studied pathways, and indicating the existence of additional layers of cellular responses.

## SUPPLEMENTARY FIGURE LEGENDS

**Figure EV1: RNP granules are assembled in response to stressors and infection in multiple cell lines.** Cell lines (**A**) HEK293, (**B**) HeLa and (**C**) HBEC were infected with Ad5 WT, **Δ**VA or **Δ**E4 at an MOI of 10, or treated with sodium arsenite (SA, 0.5 mM for 1 h) and poly(I:C) (0.5 µg/mL for 6 h), as indicated. Infected cells were fixed at 24 hpi for Ad5 WT and **Δ**VA, or at 40 hpi for **Δ**E4. Cells were stained for the granule markers G3BP1 (green) and PABPC1 (magenta), and nuclei is marked with DAPI (blue). SGs were observed in SA-treated and ΔVA-infected cells. RLBs were observed in poly(I:C)-treated and ΔE4-infected cells. No granules were observed in untreated or WT-infected cells. Scale bar = 15 µm.

**Figure EV2: Nuclear PABPC1 localization during late-stage of WT and ΔVA infections.** (**A**) Representative fields of Ad5 WT and **Δ**VA-infected A549 cells at 40 hpi, showing cells in which PABPC1 is either detected exclusively in the cytoplasm or exhibits a distinct nuclear localization. (B) The proportion of cells showing nuclear PABPC1 signal was quantified for both WT and ΔVA infections. (**C**) Nuclear PABPC1 was also quantified in **Δ**VA-infected cells which were either positive or negative for stress granules, showing similar proportions between both groups. Scale bar = 15 µm.

**Figure EV3: APEX2-based granule proteomics captures known SG proteins.** (**A**) Schematic representation of APEX2 proximity labeling, indicating selective targeting of proteins in close proximity to G3BP1 (∼20 nm radius). (**B**) Immunofluorescence in HEK293T cells expressing G3BP1-APEX2-GFP following treatment with sodium arsenite (SA, 1.0 mM for 1 h) or poly(I:C) (1.0 µg/mL for 4 h). Cells were co-stained for PABPC1 (red) and nuclei (DAPI). Scale bar = 15 µm. **(C-G)** Data normalization and reproducibility of APEX2 proteomic datasets across untreated (light grey), mock (dark grey), SA (green), and poly(I:C) (magenta) treatments for each replicate. (**C-D**) Boxplots showing log2 transformed (**C**) and median normalized APEX abundance data (**D**) for each replicate. (**E**) Bar charts showing the number of APEX proteins identified for each replicate and their intersection across replicates. (**F**) Venn diagrams showing the overlap of APEX proteins identified between replicates. (**G**) Correlation plots showing correlation coefficients for APEX abundance comparions across replicates. Correlation plots were generated using the corrplot package in R (version 0.95 built in R version 4.4.1) with correlations calculated for pairwise complete observations. (**H**) Distribution of SA APEX z-scores for all proteins identified by G3BP1-APEX proximity labeling during SA treatment (green) and proteins identified by Markmiller *et al*. (grey). The Venn diagram shows the intersection of predicted SA-induced stress granule proteins in our dataset (defined by SA z-score > 0) and stress granule proteins identified by Markmiller *et al*. The z-score threshold was selected based on the overlap between the two datasets. (**I**) Distribution of poly(I:C) APEX z-scores for all proteins identified by G3BP1-APEX proximity labeling during poly(I:C) treatment (magenta) and proteins identified by Burke *et al*. (grey). Venn diagram showing intersection of predicted poly(I:C)-induced RLB proteins in our dataset (defined by poly(I:C) z-score > 0) with proteins identified in Burke *et al*. The z-score threshold was selected based on the overlap between the two datasets.

**Figure EV4: Cellular proteins differentially recruited to SGs and RLBs.** (**A-C**) A549 cells were infected with WT Ad5, **Δ**VA and **Δ**E4 MOI 10 for 40h or treated with sodium arsenite (SA, 0.5 mM 1 h) and poly(I:C) (0.5 μg/mL for 6 h) as indicated. Panels show validation of RNA binding proteins identified by APEX analysis as being differentially recruited to SGs and RLBs. G3BP1 staining (green) was used as a marker for granule assembly, and nuclei are stained with DAPI (blue). (**A**) Staining for FMRP and Caprin1 (magenta) show the recruitment of proteins to both granules. (**B**) Staining for PRRC2C and AGO2 (magenta) indicates differential granule composition, with proteins present only in SGs induced by sodium arsenite or **Δ**VA infection and absent from RLBs. (**C**) Co-staining of G3BP1 with P-body proteins LSm14A and Dcp1b demonstrates close proximity between SGs/RLBs and P-bodies, with partial recruitment of LSm14A to both types of granules. Scale bar = 15 µm (main panels), 2 µm (cropped panels).

**Figure EV5:** RNP granule formation in G3BP1/2 KO cells during stress and viral infection. (A) A549 cells lacking expression of G3BP1 and G3BP2 (G3BP1/2 KO) were infected with Ad5 WT, **Δ**VA and **Δ**E4 MOI 10 for 24 h, or treated with sodium arsenite (SA, 1.0 mM 1 h) and poly(I:C) (0.5 μg/mL for 6 h), as indicated. Cells were stained for PABPC1 (magenta) and FXR1 (green), showing RLB assembly in response to poly(I:C) and **Δ**E4 infection. (**B**) Alternatively, G3BP1/2 KO cells were treated with SA, poly(I:C) or infected with **Δ**E4 and stained for FXR1 (magenta) and UBAP2L (green). Nuclei were stained with DAPI (blue). Scale bar = 15 µm (main panels), 2 µm (cropped panels).

**Figure EV6: Assembly of RLB-like granules during ΔE4 infection is not dependent on RNase L in U2OS cells.** (**A**) U2OS parental and RNase L KO cells (**B**) were treated with poly(I:C) (1 µg/mL for 6 h) or infected with **Δ**E4 at an MOI of 10 for 40 h. Cells were stained for G3BP1 (green) and PABPC1 (magenta), and with DAPI for nuclei (blue). Poly(I:C) induces RLB and stress granule assembly in parental and RNase L KO cells, respectively. In contrast, **Δ**E4 infection induces RNP granules resembling RLBs in both cell lines. Scale bar = 15 µm.

**Figure S7 Poly(I:C) and ΔE4 infection promote translation arrest in A549 parental and RNase L KO cells.** (**A**) A549 parental and RNase L KO cells (**B**) were treated with poly(I:C) (1 µg/mL for 6 h) or infected with **Δ**E4 (MOI 10 for 40 h). Before fixation, cells were treated with puromycin (10 µg/mL) for 10 min to measure translation. Cells were stained for puromycin incorporation (magenta) and G3BP1 (green). Nuclei were stained with DAPI (blue). Scale bar = 15 µm.

## Materials and Methods

### Cell culture

Parental cell lines were obtained from American Type Culture Collection (ATCC) and cultured at 37 °C and 5% CO_2_. A549 cells (CCL-185), HeLa cells (CCL-2), and HEK293 cells (CRL-1573) were maintained in DMEM (Gibco, 11965084) supplemented with 1% v/v sodium pyruvate (Gibco, 11360070) 10% v/v FBS (Avantor, 89510-186) and 1% v/v Pen/Strep (100 U/ml of penicillin, 100 μg/ml of streptomycin, Gibco, 15140-122). W162 cells (Vero cells containing an integrated copy of the AdV E4 region) were a kind gift from G. Ketner (Weinberg & Ketner, 1983). HBEC-KT3 cells (CRL-4051) were grown in Airway Epithelial Cell Basal Medium (PCS-300–030) supplemented with Bronchial Epithelial Cell Growth Kit (PCS-300–040) and 1% v/v Pen/Strep. Knockout A549 cell lines lacking individual dsRNA sensors (PKR, OAS1, OAS2, OAS3 or RNase L) were previously described (Li *et al*, 2016; Price *et al*, 2022a). A549 cells with double knockout for PKR and RNase L were generated using the same CRISPR-based approach described for the single knockout lines (Li *et al*, 2016; Price *et al*, 2022a). A549 cells with double knockout for G3BP1 and G3BP2 were previously described (Burke *et al*, 2024). HEK293T cells expressing G3BP1-V5-APEX2-GFP fusion protein were kindly provided by G. Yeo (Elmsaouri *et al*, 2022; Markmiller *et al*, 2018). U2OS parental and RNase L knockout cells were a gift from R. Buisson (Manjunath *et al*, 2023). All cell lines were routinely tested for mycoplasma contamination using the LookOut Mycoplasma PCR Detection Kit (Sigma-Aldrich, MP0035-1KT).

### Viral infection

Adenovirus serotype 5 (Ad5) was originally purchased from ATCC. Ad5 ΔE4 mutant *dl1004* was previously described and obtained from G. Ketner (Bridge & Ketner, 1989). Ad5 ΔVA I/II (*dl-sub720*) was previously described and obtained from C. Sullivan (Bhat & Thimmappaya, 1984).

ΔE4 virus was expanded and titered on complementing W162 Vero cells (Weinberg & Ketner, 1983). ΔVA virus was expanded and titered on A549 PKR KO cells. All viruses were purified using two sequential rounds of ultracentrifugation in cesium chloride gradients and stored in virus dilution solution (10 mM Tris-HCl pH 8.0, 100 mM NaCl, 0.1% w/v BSA, 40% v/v glycerol) at −20 °C (short term) or −80 °C (long term). Viral stock titer was determined by plaque assay, and all subsequent infections were performed at a multiplicity of infection (MOI) of 10 PFU/cell and harvested at indicated hours post infection (hpi). Cells were infected at 80-90% confluency by incubation with diluted virus in a minimal volume of low serum growth media (2%) for 2 h. After infection viral inoculum was removed and full serum growth media was replaced for the duration of the experiment.

### Drugs and treatments

Poly(I:C) (HMW, InvivoGen, tlrl-pic) were transfected at 0.5 µg/mL or 1.0 µg/mL, as described. For transfection, poly(I:C) was complexed with lipofectamine 2000 (Thermo Scientific, 11668027) in Opti-MEM (Gibco, 31985070) following manufacturer’s instructions. Sodium arsenite (Sigma-Aldrich, S7400) was used at 0.5 mM or 1.0 mM, as indicated. For the cycloheximide (CHX, Sigma-Aldrich, C4859) treatment, CHX was added at 100 µg/mL following transfection with poly(I:C), or at 24 hpi following ΔE4 infection, with cells fixed at 40 hpi. ISRIB (MedChem Express, HY-12495A) 200 nM was added to cells simultaneously to SA and poly(I:C) treatments, or at 24 hpi following ΔE4 infection, with cells fixed at 40 hpi. Puromycin (Gibco, A1113802) treatment was performed at 10 µg/mL for 10 min to assess ongoing translation.

### Antibodies and inhibitors

The following primary antibodies were used for cellular proteins: Total PKR (Abcam ab32506, WB: 1:1,000), pT446 PKR (Abcam ab32036, WB: 1:1,000), total eIF2α (Cell Signaling 5324S, WB 1:1,000), pS51 eIF2α (Sigma-Aldrich SAB5701788, WB 1:500), GAPDH (GeneTex GTX100118, WB 1:10,000), IRF3 (Thermo Fisher 51-3200, WB 1:1,000), pS386 IRF (GeneTex GTX62132, 1:500), RNase L (Santa Cruz sc-74405, 1:500), G3BP1 mouse (DB 611126, IF 1:400), G3BP1 rabbit (Proteintech 13057-2-AP, IF 1:400), PABPC1 (Abcam ab21060, IF 1:600), UBAP2L (Bethyl Laboratories A300-534A-T, IF 1:100), FXR1 (EMD Millipore 05-1529, IF 1:100), eIF4G1 (Abcam ab2609, IF 1:100), FAM120A (Sigma-Aldrich HPA019734, IF 1:500), FMRP (EMD Millipore 05-1235, IF 1:250), Caprin1 (Proteintech 15112-1-AP, IF 1:200), Ago2 (Abnova H00027161-M01, IF 1:100), PRRC2C (Thermo Scientific PA5-55936, IF 1:100), LSm14A (Proteintech 18336-1-AP, IF 1:100), Dcp1b (Cell Signaling 13233S, IF 1:100). The following antibodies against dsRNA mouse 9D5 (EMD Millipore 3361, IF 1:2) and puromycin (Sigma-Aldrich MABE343, IF 1:500) were used. For detection of viral proteins, we used anti-adenovirus 5 (against late capsid proteins, e.g. Hexon, Penton, and Fiber) (Abcam ab6982, WB 1:10,000), anti-DBP mouse (gift from D. Ornelles, Clone: B6-8, WB 1:500, IF 1:200), E4orf6 mouse RSA3 (gift from P. Hearing, WB 1:500).

### RNA isolation, qRT-PCR and Bioanalyzer

Total RNA was isolated from cells by either TRIzol extraction (Thermo Fisher) or RNeasy Micro kit (Qiagen, 74106), following manufacturer protocols. RNA was treated with RNase-free DNase I (Qiagen, 79256), either on-column or after ethanol precipitation. RNA was converted to complementary DNA (cDNA) using 1 μg of input RNA in the High-Capacity RNA-to-cDNA kit (Thermo Fisher, 4387406). Quantitative PCR was performed using the standard protocol for SYBR Green reagents (Thermo Fisher, 4367659) in a QuantStudio 7 Flex Real-Time PCR System (Applied Biosystems). All primers were used at 10 μM. All values were normalized by the ΔΔCt method by normalizing first to *GAPDH* transcripts. Ribosomal RNA integrity was assessed by running total RNA extracts on RNA chips (Agilent RNA 6000 Nano Kit, 5067-1511) using an Agilent 2100 BioAnalyzer. The following primers were used (5’-3’): *IFNB1* (Fp: CAGCATCTGCTGGTTGAAGA, Rp: CATTACCTGAAGGCCAAGGA); *IRF7* (Fp: GATCCAGTCCCAACCAAGG, Rp: TCTACTGCCCACCCGTACA); *GAPDH* (Fp: TGCACCACCAACTGCTTAGC, Rp: GGCATGGACTGTGGTCATGAG).

### Primer extension assay and RNA gel electrophoresis

Total RNA was isolated from cells by TRIzol extraction (Thermo Fisher) following manufacturer protocols. First strand synthesis buffer (Thermo Scientific) and 10 µM of 5’-IRDye800 labeled VA I or II-specific primers were added to equal amounts of RNA for each sample and denatured at 80 °C for 10 min, followed by annealing at 56 °C for 2 h. Elongation mix (100 U SuperScript III (Thermo Scientific), RNase inhibitor (NEB), 2 mM DTT (Thermo Scientific), 1 mM dNTPs (Fischer Scientific), and 10 ng/µl actinomycin D (Cayman Chemical Company) in 1X First Strand Synthesis buffer) was added to each sample and incubated at 55°C for 1 h. RNA was ethanol precipitated, resuspended in 1X RNA Loading Dye, separated on 10% acrylamide, 8 M Urea/TBE gels, and imaged on a LiCOR CLx Odyssey scanner at 800 nm. The following primers were used (5’-3’, 5’-IRDye800 labeled): VA RNA I (TTGTCTGACGTCGCACACCTG); VA RNA II (TCCGGAGGAATTTGCAAGCGG).

### Indirect immunofluorescence assays and analysis

Cells were grown on glass coverslips in 24-well plates. After indicated treatments or viral infection, cells were washed with PBS and then fixed in 4% w/v paraformaldehyde (PFA) for 10 min. Cells were permeabilized with 0.5% v/v Triton X-100 in PBS for 10 min and blocked in blocking solution (5% w/v BSA in PBS, 0.1% v/v Tween 20, 0.05% w/v sodium azide) for 1 h. Primary and secondary antibody dilutions were added to coverslips in blocking solution for 1 h, with three PBS washes in between. Nuclei were stained with 4,6-diamidino-2-phenylindole (DAPI) together with secondary antibodies. Secondary antibodies conjugated to Alexa Fluor 488, Alexa Fluor 568, or Alexa Fluor 647 (Invitrogen) against mouse or rabbit were used at a 1:500 dilution. Coverslips were mounted onto glass slides using ProLong Glass Antifade Mountant (Thermo Scientific P36984). Immunofluorescence was visualized using a Zeiss LSM 980 Confocal microscope (Cell and Developmental Microscopy Core at UPenn) and ZEN Blue v. 3.5 software. Images were processed in FIJI and assembled in Figma. Quantification of fluorescence intensity or granule-positive cells was performed from randomly acquired fields. Translational activity was measured by quantifying puromycin fluorescence in FIJI and plotted as the mean fluorescence intensity (MFI) for the whole cell area of individual cells. For ΔE4 infection, puromycin signal was quantified in cells displaying granules or nuclear PABPC1 localization. Background was calculated from cell-free regions and subtracted from intensity values.

### Fluorescence *in situ* hybridization

Cells were grown on glass coverslips followed by poly(I:C) transfection or viral infection. At indicated time points, cells were fixed in ice-cold 70% ethanol overnight at 4 °C. After fixation, coverslips were washed with PBS and treated with RQ1 RNase-free DNase (Promega, M6101) for 1 h at 37 °C, followed by two washes in PBS containing 0.1 M EDTA. Primary and secondary antibodies were incubated in PBS for 1 h skipping blocking, with three PBS washes in between. Following secondary staining, cells were fixed in 4% PFA for 10 min and washed twice in PBS. Fluorescence *in situ* hybridization (FISH) or single molecule FISH were performed for polyadenylated RNA and *GAPDH* transcripts using Stellaris ShipReady probes (Biosearch Technologies, T30-ATTO647N-1 and SMF-2019-1) and buffers. In brief, coverslips were incubated in Buffer A (1X Buffer A supplemented with formamide, SMF-WA1-60) for 5 min and then incubated overnight at 37 °C with probes diluted in hybridization buffer (SMF-HB1-10). Coverslips were then washed twice in Buffer A for 30 min (with DAPI added to the last wash), rinsed in Buffer B (SMF-WB1-20) for 5 min, and mounted on glass slides. The level of *GAPDH* transcripts was quantified as total fluorescence per cell and reported as Corrected Total Cell Fluorescence (CTCF = area × [mean intensity − mean background]). For poly(I:C) and ΔE4 conditions, *GAPDH* mRNA signal was quantified in cells displaying granules or nuclear PABPC1 localization.

### SDS-PAGE and immunoblotting

For immunoblotting analysis, cells were lysed in RIPA buffer (50 mM Tris-HCl pH 7.5, 150 mM NaCl, 1% Triton X-100, 0.5% sodium deoxycholate, 0.1% SDS) supplemented with 1% v/v Halt protease and phosphatase inhibitor cocktail (Thermo Scientific, 78446). Protein concentration was determined using a BCA assay kit (Thermo Scientific, 23227). Lysates were mixed with NuPAGE LDS Sample Buffer (4X) containing 1% v/v β-mercaptoethanol and boiled for 10 min, at 95 °C. Proteins were resolved by SDS–PAGE on NuPAGE 4-12% Bis-Tris gels (Invitrogen) using MOPS SDS running buffer (Invitrogen, NP0001). Proteins were transferred to 0.2 μm nitrocellulose membranes (Cytiva Amersham, 10600004) in transfer buffer (25 mM Tris, 192 mM glycine, 10% methanol). Membranes were blocked in 5% w/v non-fat dry milk in TBST (20 mM Tris pH 7.5, 150 mM NaCl, 0.1% Tween-20) supplemented with 0.05% sodium azide. Primary antibodies were incubated overnight at 4 °C in blocking buffer. HRP-conjugated secondary antibodies were incubated for 1h at room temperature, with three TBST washes in between. Immunoblots were developed using SuperSignal West Pico PLUS chemiluminescent substrate (Thermo Scientific, 34580) and imaged using a Syngene G-Box.

### APEX2 proximity labeling

APEX2-mediated proximity labeling was carried out essentially as described by the Yeo Lab (Markmiller *et al*, 2018; Elmsaouri *et al*, 2022). Briefly, HEK293T G3BP1-V5-APEX2-GFP cells were seeded in 10 cm plates to be confluent the following day and were either left untreated, treated with sodium arsenite (SA, 0.5 mM for 1 h), or transfected with poly(I:C) (1.0 µg/mL for 4 h) to induce granule formation. One hour prior to labeling (or at the time of SA treatment), 500 µM Biotin Tyramide (R&D Systems, 6241/25) was added to the culture media. Labeling was initiated by adding hydrogen peroxide (Sigma-Aldrich H1009) to a final concentration of 1 mM for 60 s before quenching the biotinylation reaction with Trolox (Cayman Chemical Company, 10011659) and sodium L-ascorbate (Sigma-Aldrich, A7631) at final concentrations of 5 and 10 mM, respectively. Cells were then washed once with ice-cold quenching solution and lysed in plate, on ice, in 500 µl ice-cold 8 M Urea Lysis Buffer (8 M Urea, 150 mM NaCl, 20 mM Tris-HCl pH 8.0, 5 mM Trolox, 10 mM Sodium Ascorbate, 10 mM Sodium Azide, 1X Halt Protease/Phosphatase inhibitor (Themo Scientific, 78445)). Cell lysates were sonicated and cleared by centrifugation at 13,500 x g for 10 min at 4 °C. Protein concentration was determined using the Pierce 660 nm Protein Assay (Themo Scientific, 22660), and equal amounts of protein were reduced with 10 mM TCEP (Thermo Scientific, 77720), alkylated with 15 mM iodoacetamide (Sigma-Aldrich, I1149) in the dark at room temperature for 45 min and inactivated with 10 mM DTT in the dark at room temperature for 15 min. Samples were diluted to 2 M urea by adding 3 volumes of 150 mM NaCl, 20 mM Tris-HCl pH 8.0 with protease inhibitors and quenchers. For affinity purification, 100 µl of streptavidin magnetic beads (Fischer Scientific PI88817) per sample were equilibrated with IP Wash Buffer (2 M Urea, 150 mM NaCl, 20 mM Tris-HCl pH 8.0). Samples were added to the equilibrated beads, incubated with rotation at room temperature for 2 h, and washed 6 times with IP Wash Buffer. After aspirating the final wash, the beads were resuspended in 50 mM triethylammonium bicarbonate (TEAB, Sigma-Aldrich, T7408), frozen on dry ice, and stored at −80 °C until subsequent on-bead digestion. The IP samples were subjected to on-bead digestion using endoproteinase Lys-C (Wako) at a 1:100 (w/w) enzyme-to-substrate ratio. Samples were incubated for 1 hour at 37 °C. Following Lys-C digestion, CaCl₂ was added to a final concentration of 1 mM, and sequencing-grade trypsin (Promega) was added at 500 ng per sample. Corresponding input samples for each IP were diluted to a final Urea concentration of 1 M in 50 mM TEAB. Lys-C digestion was performed as described above, followed by trypsin digestion at a 1:50 (enzyme-to-protein) ratio at 37 °C overnight with agitation. Digestions were quenched by the addition of formic acid (FA) to a final concentration of 5%, achieving a pH ≤ 3. Peptides from both IP and input samples were desalted using Poros Oligo R3 reverse-phase columns (Applied Biosystems, Thermo Scientific) packed into P200 stage tips with C18 3M plugs (3M Bioanalytical Technologies). Purified peptides were dried by lyophilization and stored at −20 °C until further analysis.

### Nanoflow liquid chromatography–tandem mass spectrometry (nLC–MS/MS)

The peptide mixture was separated using a Dionex Ultimate 3000 high-performance liquid chromatography (HPLC) system (Thermo Scientific) equipped with a two-column setup, consisting of a reversed-phase trap column (Acclaim PepMap100 C18, 5 μm, 100 Å, 300 μm i.d. × 5 mm, Thermo Scientific) and a reversed-phase analytical column (35 cm, 75 μm i.d., 360 μm o.d., packed with Pur C18AQ, 2.4 μm; Dr. Maisch). Loading buffer was 0.1% trifluoroacetic acid (Merck Millipore) in water. Buffer A was 0.1% formic acid, and Buffer B was 80% acetonitrile (ACN) + 0.1% formic acid. The HPLC was coupled online with a Q-Exactive-HF mass spectrometer (Thermo Scientific). Peptides were eluted using a 140 min ACN gradient (90 minute 5%–25% ACN gradient, followed by a 30 minute 25%–45% ACN gradient, a 10 minute 45%-95% gradient, with a final 10-minute isocratic step at 5% ACN) at a flow rate of 300 nl/min for IP and input samples. The MS instrument was controlled by Xcalibur software (Thermo Fisher Scientific). Samples were batch-randomized to account for instrument variation. The data dependent acquisition (DDA) MS method was designed with the MS1 having a window of 400–1000 m/z, AGC target of 1e6 and maximum inject time (MIT) of 75 ms with the MS2 having automated windows, AGC target of 100% and MIT of 75 ms. Fragmentation was performed with high-energy collisional dissociation (HCD) using normalized collision energies (NCE) of 27%. The selection for ions were charges 2–8, minimum peak intensity of 1e4, and a 3 s maximum cycle time.

### Proteomics enrichment analysis

The raw mass spectrometer files were processed for protein identification using the Proteome Discoverer (v2.4, Thermo Scientific) and the Sequest HT algorithm with a peptide mass tolerance of 10 ppm, fragment m/z tolerance of 0.02 Da, and a false discovery rate (FDR) of 1% for proteins and peptides. Quantification was performed using a label-free approach using the “Precursor ions quantifier” node, and peptide abundances were rolled up into protein abundance using the summed abundance algorithm using only unique or razor peptides. All peak lists were searched against the UniProtKB/Swiss-Prot database of Human sequences (9606; downloaded November 2022) using the parameters as follows: enzyme, trypsin; maximum missed cleavages, 2; fixed modification, carbamidomethylation (C); variable modifications, oxidation (M), protein N-terminus acetylation. All subsequent protein-level analysis of streptavidin-enriched APEX2-labeled proteins (APEX) and total lysate control (Input) abundance quantification data was performed using custom R scripts. Proteins were filtered to include only those identified by at least 1 unique peptide and with peptide q-value < 0.01. APEX and Input abundance quantification values within each sample were transformed to log2 values and normalized by the sample median to account for technical variation in the abundances across samples. APEX and Input abundance means for each protein, in each condition, were obtained by calculating the arithmetic average of the abundance quantification values across the three biological replicates of each condition. Biological replicates for which abundance quantification values were not obtained were removed from the calculation of the mean. Relative protein APEX and Input abundances within each condition were obtained by calculating the statistical z-scores based on the average and standard deviation within the average abundance quantifications of the respective condition. APEX abundances were normalized by the respective input abundances for each replicate of all infections and treatments. Log2 fold changes of APEX abundances for compared conditions were obtained by comparing the average normalized quantification of the respective conditions. Log2 fold changes were imputed for cases in which the protein was identified in 3/3 replicates in one condition and 0/3 replicates in the compared condition. In these cases, the log2 fold change was defined as the minimum or maximum of the fold changes within the respective condition. Statistical p-values associated with the APEX abundance log2 fold changes were calculated using unpaired, two-sided, student’s t-tests comparing the three replicate normalized quantifications of each compared condition. P-values were calculated only for comparisons in which each condition had quantifications in at least 2 of 3 biological replicates. When comparing APEX abundances across conditions, statistically enriched proteins were defined as those proteins that were identified in 3/3 biological replicates in one condition and 0/3 replicates in the compared condition, or identified in 3/3 biological replicates in one condition and exhibiting a log2 fold change > 1 and p-value < 0.05 versus the compared condition.

### Reproducibility and statistics

Experimental observations were confirmed in at least three independent biological replicates. Statistical analyses were performed using GraphPad Prism (version 10.6.0). Statistical tests, error bars, n values, and p-values are specified in the corresponding figure legends.

## Data availability

Mass spectrometry proteomics data will be deposited to ProteomeXchange Consortium (http://proteomecentral.proteomexchange.org) via the PRIDE partner repository (Vizcaíno *et al*, 2013). Alternativelly, unprocessed data will be made available upon request.

## Acknowledgments

We thank members of the Weitzman Lab for insightful discussions and input. We are grateful to A. Berk, P. Hearing, G. Ketner, A. Levine, R. Buisson, C. Sullivan, G. Yeo and D. Ornelles for generous gifts of viruses, cells, antibodies, and reagents. We thank the University of Pennsylvania Cell and Developmental Biology Microscopy Core for imaging assistance.

## Author Contributions

R.T.S., O.B.S., A.M.P. and M.D.W. conceived the project and designed experiments; S.R.W. and J.M.B. provided additional input into study design; R.T.S., O.B.S., H.C. and S.B. performed experiments; K.K. and R.L. prepared and loaded samples for mass spectrometry; J.M.D. performed computational proteomic analyses; A.R.N.A., N.K. and E.L.A. assisted with immunofluorescence acquisition and analysis; Y.L. and N.A.P. generated KO cell lines; R.T.S. and O.B.S. analyzed all additional data; R.T.S., O.B.S., A.M.P. and M.D.W. wrote the manuscript; all authors read, edited, and approved the final paper.

